# Neolithic introgression of *IL23R*-related protection against chronic inflammatory bowel diseases in modern Europeans

**DOI:** 10.1101/2024.08.06.606840

**Authors:** Ben Krause-Kyora, Nicolas Antonio da Silva, Elif Kaplan, Daniel Kolbe, Archaeological Civilization Disease Consortium (ACDC), Inken Wohlers, Hauke Busch, David Ellinghaus, Amke Caliebe, Efe Sezgin, Almut Nebel, Stefan Schreiber

## Abstract

**Background:** The hypomorphic variant rs11209026-A in the *IL23R* gene provides significant protection against immune-related diseases in Europeans, notably inflammatory bowel disease (IBD). Today, the A-allele occurs with an average frequency of 5% in Europe.

**Methods:** This study comprised 251 ancient genomes from Europe spanning over 14,000 years. In these samples, the investigation focused on admixture informed analyses and selection scans of rs11209026-A and its haplotypes.

**Findings:** rs11209026-A was found at high frequencies in Anatolian Farmers (AF, 18%) where it was likely under weak positive selection. AF later introduced the allele into the ancient European gene-pool. Subsequent admixture caused its frequency to decrease and formed the current southwest-to-northeast allele frequency cline in Europe. The geographic distribution of rs11209026-A may influence the gradient in IBD incidence rates that are highest in northern and eastern Europe.

**Interpretation:** Given the dramatic changes from hunting and gathering to agriculture during the Neolithic, AF might have been exposed to selective pressures from a pro-inflammatory lifestyle and diet. Therefore, the protective A-allele may have increased survival by reducing intestinal inflammation and microbiome dysbiosis. The adaptively evolved function of the variant likely contributes to the high efficacy and low side-effects of modern IL-23 neutralization therapies for chronic inflammatory diseases. This study highlights how evolutionary informed research can provide promising targets for new therapeutic strategies.

**Funding:** Deutsche Forschungsgemeinschaft (DFG German Research Foundation) under Germany’s Excellence Strategy – EXC 2167 390884018 and EXC 2150 390870439.

## Introduction

Inflammatory bowel disease (IBD) with its two main sub-phenotypes Crohn’s disease (CD) and ulcerative colitis (UC) is archetypical for many chronic inflammatory disorders characterized by immune dysfunction, cellular inflammation and a compromised (intestinal) barrier. The rise in incidences in IBD and many chronic inflammatory disorders is attributed to a complex interplay between genetic risk factors and environmental challenges (e.g., nutrition, hygiene, antibiotic use) that are inherent to the lifestyle of populations in industrialized countries^1,2^. The genetic risk architecture of IBD comprises more than 230 lead variants and loci that show strong overlapping associations between IBD, psoriasis and other chronic inflammatory diseases^3,4,5^. While for most identified loci the pathophysiological significance remains unexplored, some association signals have been resolved to the functional level. These include variants in the IL-23 pathway, particularly in the IL-23 receptor gene *IL23R* where protective disease associated variants are functionally hypomorphic^6,7^.

IL-23 is a key cytokine in chronic inflammation. Excessive activation of the IL-23 receptor promotes the production of inflammatory mediators including IL-17, IL-22, granulocyte-macrophage colony-stimulating factor and tumor necrosis factor^8^. The function of the IL-23 receptor is modulated by natural genetic variation, for instance, at the single-nucleotide variant (SNV) rs11209026-G/A (R381Q) in the coding sequence of *IL23R.* The derived (and minor) A-allele has shown strong protective effects against immune-related diseases in European populations (e.g., CD, UC, ankylosing spondylitis, rheumatoid arthritis, psoriasis)^4,9–17^. In addition, it has been associated with increased diversity of the intestinal microbiome as well as with the rise of beneficial microbial components^18^. Functional studies have shown that the A-allele lowers stability of the IL-23R protein, thereby diminishing receptor expression at the cell surface and STAT3/STAT4 activation in response to IL-23^7^.

This downregulation leads to decreased secretion of pro-inflammatory cytokines such as IL-17 and IL-22, explaining the protective effect of the R381Q variant in IBD and other inflammatory and autoimmune disorders^19^. Therapeutic applications that mimic this effect (i.e., reduction of IL-23R by neutralization of IL-23) lead to significantly lower levels of inflammation with breakthrough clinical efficacy levels in psoriasis, CD and UC^20–26^. It is highly surprising that the high efficacy of such significant interventions is associated with only minor side-effects.

Because of the importance of the *IL23R* variant rs11209026-A in the pathophysiology and the therapeutic targeting of chronic inflammation, we have reconstructed here its evolutionary history in Europe. To this end, we have analysed genetic information on the allele and the haplotypes on which it segregates from modern and (pre)historical populations dating back up to 14,000 years. We hypothesize that the A allele was introduced into the modern gene-pool at high frequency by the first Neolithic farmers and that its current clinical frequency distribution in Europe, which parallels the incidence and prevalence gradient for IBD, is due to admixture.

## Results

### Variant rs11209026-A in modern populations

The variant rs11209026-A is found in European populations with an average frequency of approximately 5% (ranging from 0 to 21%, Supplementary Table S1). Visualising the frequencies on a geographical map of Europe and neighbouring regions (Fig. 1) reveals an increasing gradient from north to south and east to west.

**Figure 1:**
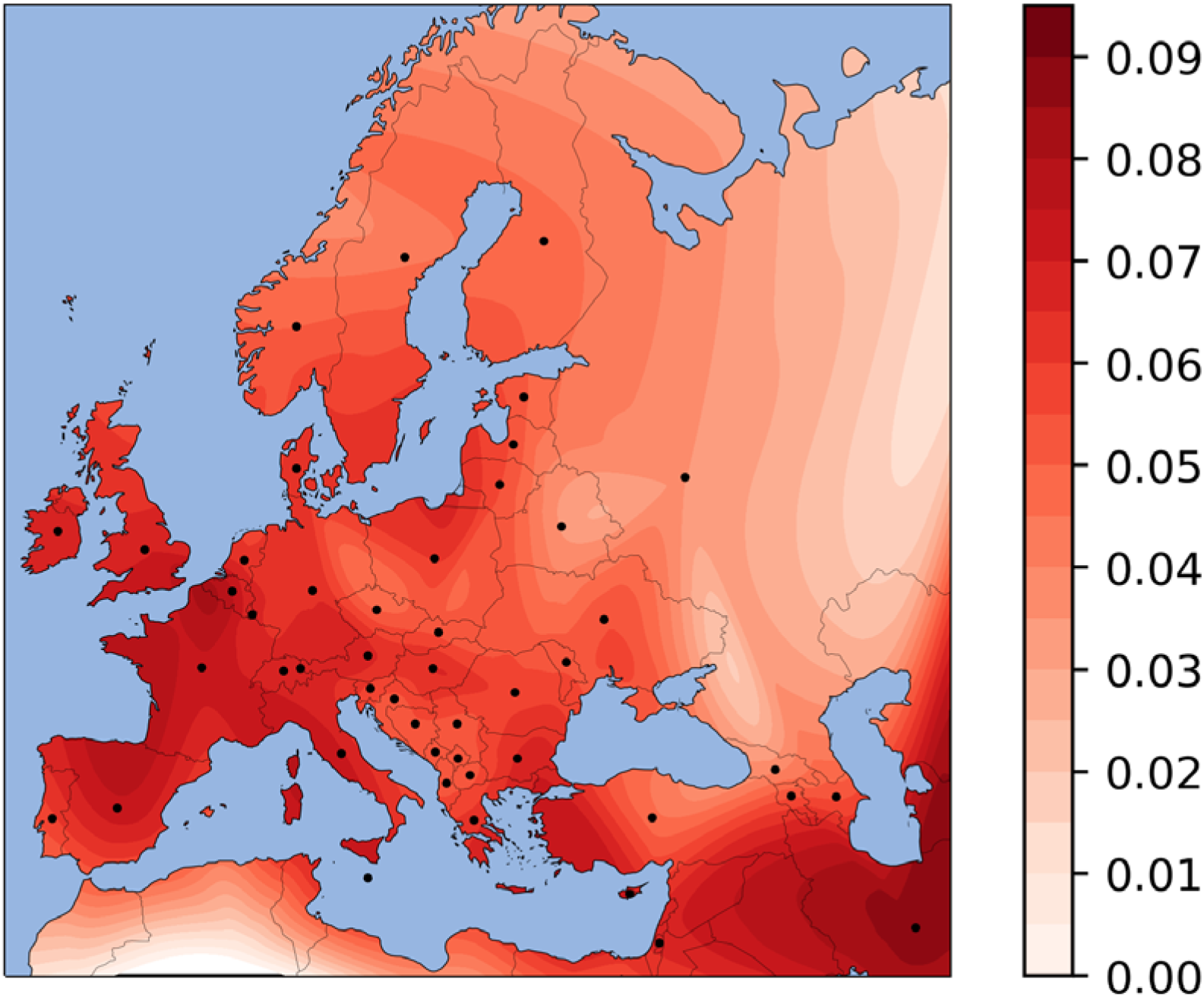
Map of rs11209026-A frequencies based on data from modern West-Eurasians (Supplementary Table 1). Colour corresponds to allele frequency according to the key bar on the right. The black dots represent the countries from which frequencies were used for the interpolation. The map shows an increasing north-south and east-west cline.

Previous studies have shown that rs11209026 lies in a larger linkage disequilibrium (LD) block on chromosome 1p31^10^. We therefore investigated in more detail the LD structure around this SNV in central Europeans (CEU). Our results revealed that rs11209026 is situated in a 42.9 kb LD-block between two recombination hotspots that does not extend into neighbouring genes (described by UCSC genome browser, Supplementary Figure S1). To analyse the haplotype structures and the complexity of this LD block, we have calculated furcation trees around rs11209026 in CEU (Supplementary Figure S2). Notably, the A-allele is present on only a few haplotypes, indicating a relatively recent origin and a short time of diversification of the A-haplotypes.

### Variant rs11209026-A in ancient populations

To gain insights into the history of rs11209026-A, we traced the frequency of the allele over time using a paleogenetic approach. For this purpose, we extracted 251 genotypes from ancient populations (Supplementary Figure S3). Of these, 66 were generated by us from Neolithic samples (Supplementary Table S2)^27,28^. The compendium provided us with a high temporal resolution going back almost 14,000 years. The final panel comprised 251 genotypes from seven populations, which differed not only in terms of time, location and subsistence, but also in their population genetic composition: Western Hunter-Gatherers (WHG), Anatolian Farmers (AF), Early Farmers (EF) and Late Farmers (LF) from the Neolithic period, Bronze Age and medieval populations from Europe as well as pastoralists from the Eurasian Steppe (Supplementary Figure S3). In addition, we used the CEU sample as a modern reference. Based on these data, we constructed a frequency trajectory over time (Fig. 2A). Interestingly, the A allele of rs11209026 was absent in the examined WHG and Steppe populations. It first appeared among AF (18.2%) and was also present in the first farmers of Europe, EF (14.3%) and LF (12.6%). After the Neolithic period its frequency decreased until it reached the present level of about 5%.

**Figure 2:**
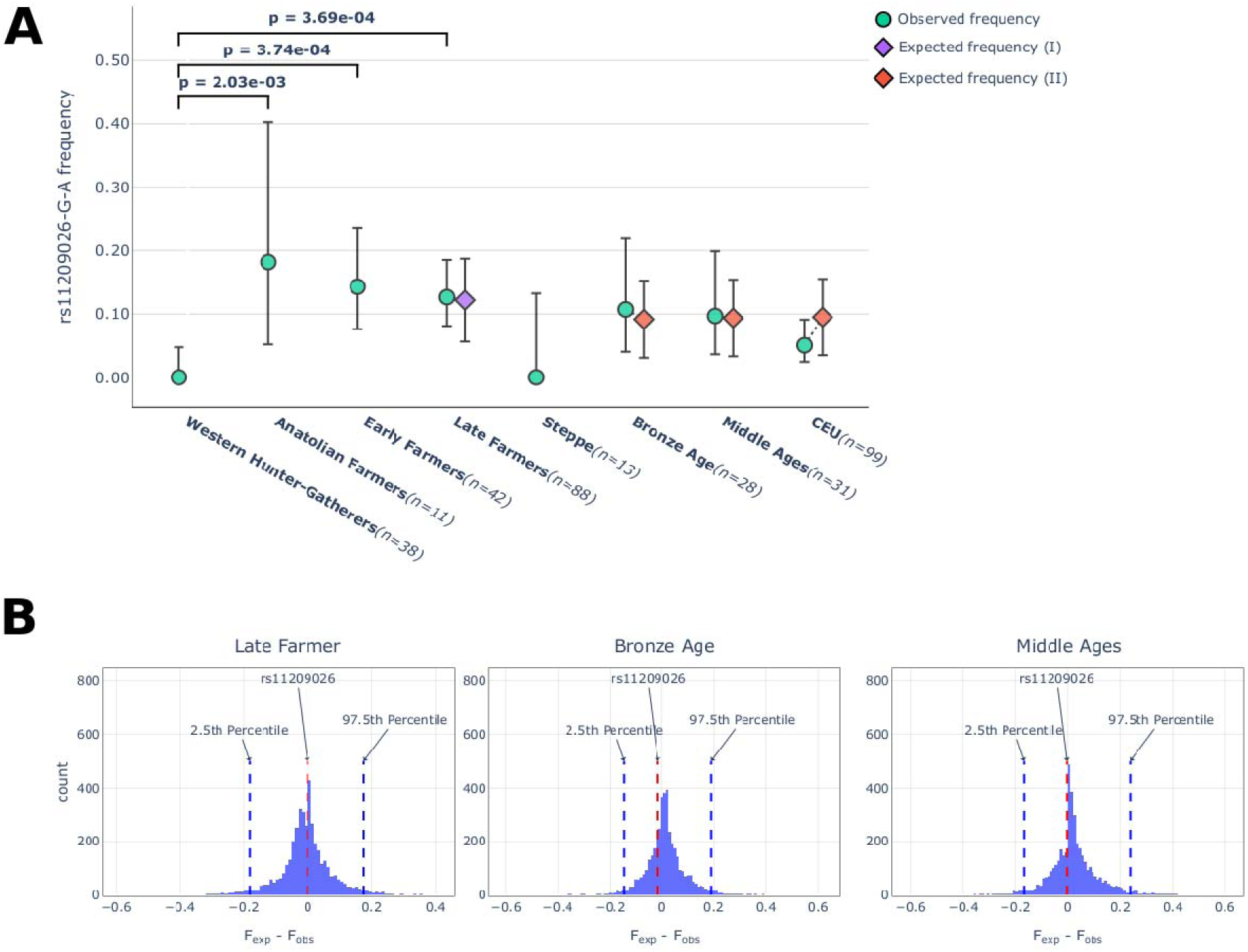
**A** Frequency of the rs11209026-A allele in seven ancient populations and a modern European reference (CEU). An expected frequency is shown next to each population resulting from one of two past major demographic events, I: admixture of western hunter-gatherers with early farmers (purple diamond); II: admixture of late farmers with Steppe herders (red diamond). The expected frequency of an offspring population was calculated using the observed frequency of its parental populations and the respective admixture component of these parental populations in the offspring population. A Fisher’s exact test was used to assess whether there were significant differences between the observed frequencies of ancient populations. For clarity, only the p-values for three population pairs (WHG vs. AF, EF, LF) are shown. Errors bars indicate the 95% CI in observed frequencies. Errors bars on the expected frequencies indicate the expected frequency range. **B** Distribution of the distances between expected allele frequencies based on admixture (F_exp_) and observed frequencies (F_obs_) of variants located in neutral regions of the genome.

### Admixture-informed analyses in ancient populations

Next, we investigated whether the decreasing allele frequency trend observed from EF to modern-day populations may have resulted from admixture. For this approach, we computed the ancestral genetic components for populations that emerged from the two major demographic events in Europe: first, the admixture of EF with WHG giving rise to LF (4500-3500 BCE); second, the massive introgression of pastoralists from eastern Steppe areas (2700-2500 BCE). We then calculated the rs11209026-A expected frequencies for these derived populations using the observed allele frequencies and genetic proportions of ancestral populations (e.g., WHG and EF as ancestral populations for LF) and assessed whether their expected and observed frequencies significantly differentiated. The observed allele frequency in LF did not significantly differentiate from the expected frequencies and was therefore most likely the result of admixture between WHG and EF (Fig. 2A). This was also the case for populations that emerged after the Steppe herders had migrated to Europe. In addition to simply comparing frequencies by taking the admixture proportions into account, we also performed a test based on the empirical distribution of the difference between expected and observed frequencies in neutral regions. This test does not only consider admixture but also genetic drift and other neutral evolutionary processes like population size variation. The frequency differences for rs11209026-A were well in the center of the empirical distribution, thus being compatible with neutral evolution under admixture (Fig. 2B). Taken together, the decreasing trend in allele frequency from EF onwards can mainly be explained by admixture.

### Selection analyses in farmers

One striking result was the significant frequency differences between WHG and each of the three farming populations, i.e., AF, EF and LF (p-value < 2 × 10^−3^, Fig. 2B), which may be due to natural selection. To identify possible selection scenarios and rule out random genetic drift, we applied numerous tests (i.e., calculations of extended haplotype homozygosity (EHH), F_ST_, Hardy-Weinberg equilibrium (HWE), Tajima’s D, Fu & Li’s D and Fu & Li’s F) in AF, EF and LF on imputed data as well as in modern CEU. Imputation is a useful approach in ancient genomics to increase the sample size and improve data quality^29,30^. We validated the imputation process in our dataset and confirmed its high accuracy with common variants, ensuring the reliability of results from haplotype-based analyses, such as EHH (Supplementary Figure S4). The EHH analysis showed that the A-allele was present in AF, albeit without an extended haplotype structure. Extended haplotypes around the A-allele first became visible in EF and were preserved in LF and CEU (Fig. 3). The presence of extended haplotypes in EF may indicate positive selection in AF. To further support the scenario of positive selection, we computed F_ST_ values for SNV/gene as well as chromosome-wide levels. The analysis showed significant frequency differences for rs11209026-A, but not the surrounding chromosomal region, between WHG and EF (Supplementary Table S3, Supplementary Figure S5). This result indicates selection on rs11209026-A. If the effect had been caused by drift, we would expect to see the same frequency differences in the SNV, the gene or the whole chromosome. However, this is not the case. Next, we calculated HWE and Tajima’s D for all three farming populations to test for deviation from neutral expectations, which could indicate strong selection (Supplementary Table S4). However, our results for rs11209026 showed no signals in any of the analysed populations, suggesting that the SNV was exposed to no (neutral expectations) or only weak selective pressures. If weak selection had indeed played a role, it would probably have mainly affected the functional A-allele carriers. Therefore, we performed haplotype-specific Tajima’s D, Fu & Li’s D, and Fu & Li’s F tests as described previously in modern populations (Table 1)^31,32^. All analyses consistently revealed statistically significant values in EF and LF, but only for the protective haplotypes with the A allele and not for the risk haplotypes with the G allele (due to small sample size in AF, the results did not reach statistical significance). All our findings (from EHH tests and haplotype-specific analyses) are consistent with weak positive selection acting on the A-haplotypes. Another interpretation of the negative Tajima’s D scores could be recent population growth. However, in this scenario, both haplotypes should have been affected, which was not the case as the risk G-haplotypes yielded neutral values throughout the analyses (Table 1).

**Figure 3:**
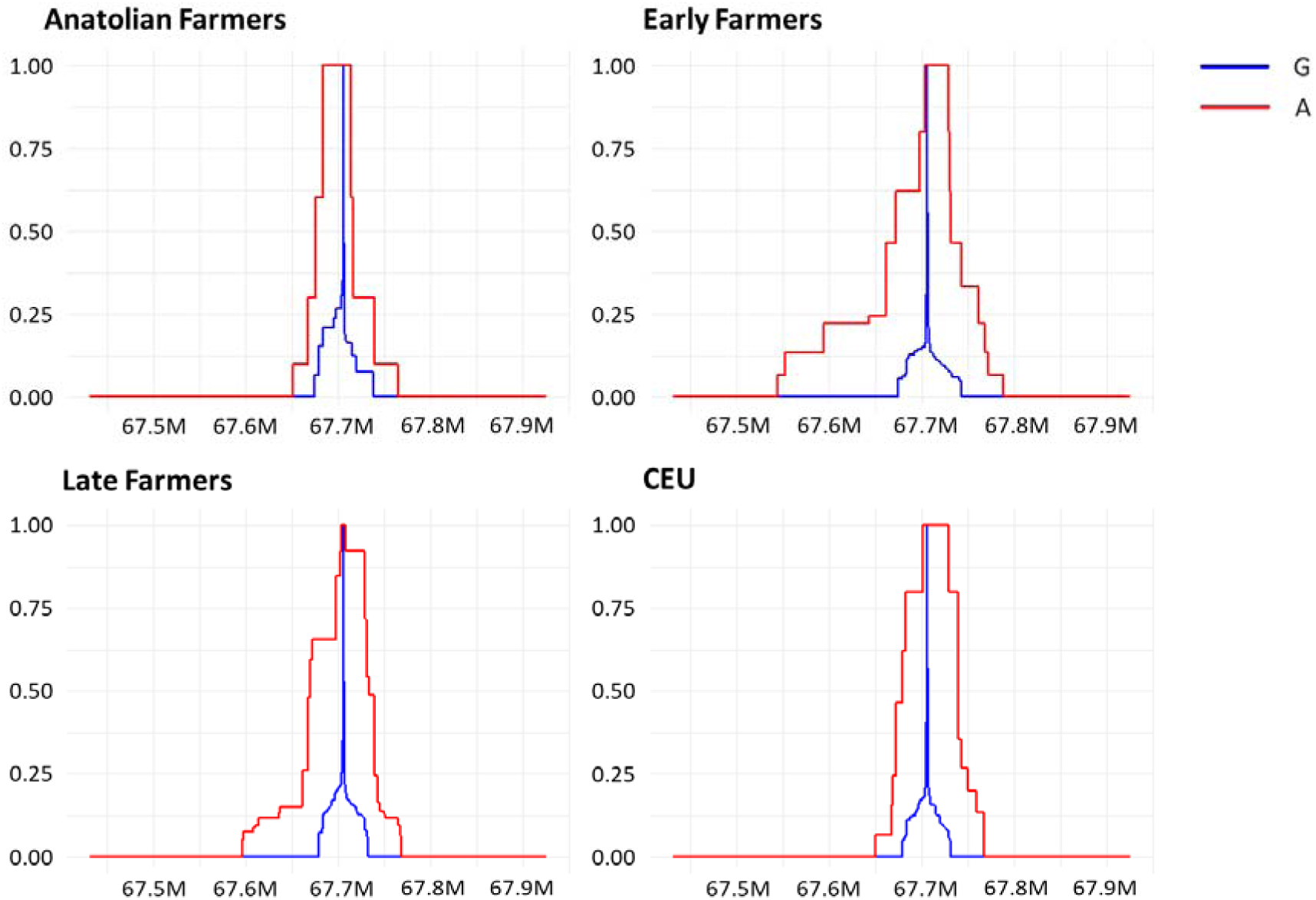
EHH plot for rs11209026 in AF, EF, LF and CEU populations. The x-axis represents the coordinates around the SNV while the y-axis shows the homozygosity scale ranging from 0 (no haplotype homozygosity) to 1 (complete haplotype homozygosity). In EF, LF and CEU the extended homozygosity is seen for the haplotype with the derived A-allele.

**Table 1:** Tajima’s D, Fu & Li’s D, and Fu & Li’s F test scores for rs11209026-A or -G haplotypes.

| Haplotype | Population (N) | Tajima's D | Fu and Li's D | Fu and Li's F |
| --- | --- | --- | --- | --- |
| **A-haplotypes** | Early Farmers (10) | -1.92* | -2.25** | -2.44** |
|  | Late Farmers (39) | -2.45*** | -4.70** | -4.65** |
|  | CEU (10) | -1.92* | -2.25** | -2.44** |
| **G-haplotypes** | Early Farmers (50) | 1 | 0.3 | 0.67 |
|  | Late Farmers (187) | 0.54 | -1.53 | -0.66 |
|  | CEU (188) | 0.32 | -2.01 | -1.07 |

Due to the low number of samples, these tests could not be performed on the AF population.
The window size is 42.9 kb.
N: Number of haplotype sequences analyzed
The values for the G-haplotypes were all non-significant.
\*, $P < 0.05$
\*\*, $P < 0.02$
\*\*\*, $P < 0.01$

The HWE results (whole CEU dataset) for a larger region surrounding *IL23R* (∼500 kb) did not show any signs of drift which could potentially have produced false-positive selection signals. To rule out possible haplotype-specific drift, we have performed Tajima’s D tests on the A-allele carriers for the same 500-kb region (Supplementary Figure S6) (with a window size of 40kb and step size of 100bp). The results showed only significant negative Tajima’s scores for the A-haplotype LD block with the extended haplotype structure (EHH).

## Discussion

In this study, we investigated the protective *IL23R* allele rs11209026-A in 251 ancient population samples. A striking finding was the statistically significant differences in allele frequencies between WHG and the three farming populations. Our detailed selection analyses on paleogenetic data (rather than simulations) suggest that these differences can most likely be explained by weak positive selection on the A-allele and its haplotypes in farmers rather than by random genetic drift or demographic processes such as population expansions. The oldest population in which we detected this signal were EF (the available data on AF were not sufficient for the analyses), indicating that the selection must have started before the formation of this group. We observed a decreasing A-allele frequency trend from the first Neolithic farmers in Europe (14.3%) to present-day populations (∼5%) which can mainly be explained by the two major admixture events known to have shaped the current gene pool^33^.

About 12,000 years ago hunter-gatherers in the Near East and Anatolia transitioned to a farming lifestyle, thus initiating the Neolithic transformation^34^. Around 5,000 years later the Anatolian Farmers (AF) migrated westwards to southern and central Europe. Along their way, they experienced little admixture with WHG, giving rise to the central European EF population^35^. Against the background of this information, it is interesting to evaluate the genetic results. While rs11209026-A was found to be completely absent in WHG, it was present in all three studied farming populations at appreciable frequencies, far above those in today’s Europeans. The highest frequency of the allele in the analysed dataset (18%) was detected in AF, in European farmers (EF, LF) it was slightly lower due to admixture with WHG (see below). Our EHH results indicate that the mutation at rs11209026 leading to the A-allele/-haplotypes likely occurred in AF. Another hypothesis could be that the mutation already arose in the preceding hunter-gatherer populations in Anatolia which are estimated to have formed around 14,000 years ago^35^. Unfortunately, so far there is no genomic data available from this group. However, the A-allele is also absent in the few available samples of Eastern Hunter-Gatherers (EHG, n=1) and Caucasus Hunter-Gatherers (CHG, n=2) covered at rs11209026^36–38^. The Steppe population, whose genomes mainly consist of an even mixture of EHG and CHG ancestry, lacked the allele as well (Fig. 2A)^39^. In view of these observations, it is likely that rs11209026-A did not arise in the common ancestors of all hunter-gatherers (around 25,600 years ago)^35^. The high frequency of the allele in AF without the corresponding extended haplotype structure could mean that it had a relatively recent origin (i.e., in the last 14,000 years).

Which events could have led to the high frequency of the *IL23R* variant rs11209026-A in AF? AF are reported to have lived in isolated groups with a small effective population size and to have been exposed to several bottlenecks over a few hundred years, which could have favored drift^32^. However, given the dramatic changes in lifestyle from hunting and gathering to agriculture, AF might also have been subject to selective pressures. Skeletal remains from AF exhibited lesions and shorter stature (compared to hunter-gatherers), both of which suggest malnutrition and/or famine^40^. These in turn may cause microbiome dysbiosis and intestinal inflammation^41,42^. It has also been hypothesized that the early farmers were confronted with a high load of various infectious and non-infectious inflammagens^43^. The farming lifestyle was sedentary and relied on the cultivation of plants and the domestication of animals. The latter involved also co-housing with domesticates which must have resulted in contact with animal excreta/microbiome and possibly zoonoses^44–46^. Relative to hunter-gatherers, the diet of the first farmers is thought to have been rather monotonous, unbalanced and rich in carbohydrates, dairy and alcohol^44,47^. Given such a pro-inflammatory environment and lifestyle, it is conceivable that the protective A-allele increased survival chances by reducing inflammatory responses in general and by dampening intestinal inflammation and dysbiosis in particular. By migrating to Europe, AF later introduced the A-haplotypes into the ancient European gene-pool in which they have been maintained ever since despite major admixture events.

After indigenous WHG and incoming EF had coexisted for nearly 2,000 years in central Europe, the WHG communities died out or were absorbed into the EF population and adopted the agricultural lifestyle^27^. The latter likely resulted in a decrease in frequencies of the protective rs11209026-A allele in later central European farming populations, here referred to as LF. A further decline is due to admixture when pastoralists from the Pontic–Caspian Steppe migrated to eastern and central Europe in large numbers during the Late Neolithic and Early Bronze Age^48^. The Steppe herders, who had large EHG/CHG ancestry proportions, also lacked rs11209026-A, which led to a further decrease in the allele in Europe. Between the Bronze Age and modern times (including the Middle Ages) we only observe small changes in frequency. Our admixture-informed analyses have shown that the overall decrease in frequency since the European Neolithic is mainly due to admixture events rather than negative selection as suggested previously^49^.

The inflow of the Steppe population reached central Europe from northern and eastern routes but did not penetrate deeply into the south and west of the continent. Frequencies of the protective *IL23R* allele rs11209026-A in today’s populations show a southwest-to-northeast gradient (Fig. 1) with the south and the west of Europe having higher frequencies, correlating with more EF ancestry. This may contribute to the well-described gradient in incidence rates for IBD with the highest incidence and lifetime prevalence seen in northern and eastern Europe^50,51^.

Our findings shed light on the potential health benefits of reducing IL-23/IL-23R activity, which may have led to increased survival in the early farmers in Anatolia and their descendants in Europe by reducing inflammatory responses, especially by dampening intestinal inflammation and dysbiosis. This genetic predisposition results in today’s world in a protection against inflammatory diseases of the intestine, joints and skin.

Anti-IL23 therapies have breakthrough efficacy in several chronic inflammatory diseases in the modern population. This unprecedented efficacy is paired with an extremely low side-effect profile without clinically relevant immune suppression, which corresponds to the positive health profile of individuals with hypomorphic *IL23R* function. While the anti-inflammatory effectiveness of anti-cytokine therapies (e.g., anti-TNF, anti-IL-1beta, anti-IL-6R) is usually paralleled by immune suppressive side-effects, it is clinically surprising that blockade of IL-23, which exerts strong anti-inflammatory efficacy in several immune mediated diseases, does not lead to opportunistic infections or other immune suppressive complications in clinical trials and real-world observations. Anti-IL-23 therapies could therefore be an example of how evolutionary principles can guide pharmaceutical developments resulting in desirable benefit/risk profiles. Future clinical development of anti-IL23 strategies that include small molecules and orally administered peptide drugs may target early intervention in patients with chronic inflammatory disease or may even examine pre-emptive or preventive effects in high-risk individuals^52^.

## Conclusion

Our results show that the protective *IL23R* allele rs11209026-A was introduced into the central European gene-pool through AF. The adaptively evolved benefit of the protective variant in early farmers may explain the low side-effects of today’s pharmaceutical interventions that mimic its effect in therapy of chronic inflammatory diseases. Therefore, evolutionary informed interventions could be a promising approach to develop new therapies.

## Materials and Methods

### Frequency map of rs11209026-A in modern populations

We compiled rs11209026-A frequency data from 33 modern populations across West-Eurasia^4,50,53–55^. Countries with small sample sizes (<30) or unavailable data had the allele frequency inferred from the average of the countries in the region for which data was available (Supplementary Table S1). An interpolated allele frequency map was then generated using SciPy^56^.

### LD and haplotype analysis in modern Europeans

We used LDBlockShow to calculate the LD structure surrounding the *IL23R* region in the central Europeans (CEU) from the 1000 Genomes Project^53,57^. The analysis was performed using only bi-allelic sites, a 0.05 minor allele frequency cut-off and 0.25 maximum ratio of missing alleles. The blocks were estimated using Gabriel’s definition^58^. The bifurcation tree of the LD block was prepared by using the “rehh” package version 3.2.2^59^. Bifurcation trees help visualize haplotype patterns and their relations.

### Ancient DNA data

Sequencing data (FASTQ) or mapping data (BAM) of individuals listed in the Allen Ancient DNA Resource (AADR version V50.0.p1) was downloaded^60^. From the whole compendium, we selected 185 individuals that had the SNV rs11209026 covered. These samples are from West-Eurasia (n=161), the Near East (n=11) and the Russian Steppe (n=13). We also included data from 66 Neolithic samples (Supplementary Table S3) that have not yet been incorporated into the AADR^27,28^. This resulted in a dataset containing 251 samples in total. Based on archaeological and population genetic criteria, the samples were grouped in seven populations (Supplementary Figure S3). Sequencing data was mapped and processed as described below.

### Processing and mapping of sequencing reads

Sequencing data was preprocessed as described elsewhere^27^ and subsequently mapped to the human genome build hg19 using BWA^61^ with the reduced mapping stringency parameter “-n 0.01”. Authenticity of the ancient DNA was verified as described before^27^. To reduce genotyping errors arising from the ends of the reads, we estimated deamination patterns with DamageProfiler v1.1^62^ and trimmed the ends of the reads with bamUtils v1.0.15^63^ to achieve a terminal damage rate below 5%. In addition, the mapped data was filtered by quality, keeping reads with mapping quality and base alignment quality greater than 20.

### rs11209026 genotyping and calculation of allele frequencies in ancient individuals

We used BCFtools v1.12 to call diploid instead of pseudo-haploid genotypes to generate more precise allele frequencies of rs11209026^64^. The following filtering criteria were used for quality control: MQ>20; BQ>30; DP ≥ 3. Existing BQ tags were ignored and BQ was recalculated on the fly (-E). The mapping quality of reads was adjusted to account for excessive mismatches expected for ancient DNA (--adjust-MQ 50). In total, we analysed a sample of 251 individuals for the SNV.

### Admixture-informed analysis

To test the influence of admixture on the rs11209026-A allele frequency trajectory (Fig. 2A), we performed an admixture-informed analysis on the 251 diploid datasets. The admixture components for each population were calculated using^65^. Two admixture events were taken into consideration. 1) WHG and EF were used as parental populations for the LF and 2) LF and Steppe herders for all populations emerging from the Bronze Age onwards. For each offspring population with the parental populations x and y, the lower and upper bounds of the expected frequency range were calculated as follows:

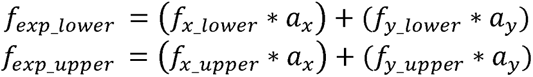

The terms *f_(x,y)_lower_* and *f_(x,y)_upper_* represent the lower and upper bounds of the 95% confidence interval of the observed frequency of either x or y. a denotes the corresponding admixture component of the parental population in the offspring population. To assess whether the observed frequency of the offspring population significantly deviates from this range, two one-sided binomial tests were employed, which utilized the *f_exp_lower_* and *f_exp_upper_* as expected frequencies, respectively. The lowest p-value from both tests was kept. Typically, this yielded a p < 0.05 if the 95% confidence interval of the observed frequency of the offspring population did not overlap with the expected frequency range calculated above.

Since the above approach is rather simplistic, as it assumes instantaneous admixture and does not account for genetic drift and population size variation, we additionally performed a more sophisticated statistical test. The corresponding test statistic is the difference between expected (F_exp_) and observed (F_obs_) frequency of the variant to be investigated (here rs11209026-A). The distribution of the test statistic under the null hypothesis is derived empirically by calculating F_exp_-F_obs_ on neutral regions of the genome compared with those of rs11209026-A. The 2.5% and 97.5% percentiles of this distribution will serve as significance thresholds. Thus, variants with F_exp_-F_obs_ values lower or higher than these thresholds are significantly deviating from neutral evolution. This novel test has the advantage that all variations in allele frequencies due to neutral evolution such as admixture, bottlenecks, genetic drift etc. are mapped in the empirical distribution. Only variants showing large differences in allele frequencies from evolutionary processes acting on putatively neutral regions will reach significance. Analogous to Gazave et al.^66^, we estimated putatively neutral regions of the genome by excluding coding or potentially coding sequences, along with an additional 100 kb (approximately 0.1 cM) buffer around these sequences. Regions containing segmental duplications, copy number variants, conserved elements, CpG islands and repetitive elements were also removed. Additionally, we excluded the centromere and 40-kb regions at the chromosome extremities (sources for all these features are available in Supplementary Table S5). Remaining genomic regions shorter than 1 kb were filtered out. The resulting set of putatively neutral regions totaling around 107 Mb spread across 972 regions of the genome. We then generated genotype calls and allele frequencies (F_exp_ and F_obs_), as described above, for 358,755 bi-allelic records within the set of neutral regions that were covered in our dataset of ancient samples. Additional filtering with plink v1.9^67,68^ for individual missingness (--mind 0.5) and minor allele frequency (--maf 0.01) yielded a final dataset containing 171 individuals and 4094 variants. This final dataset of putatively neutral regions was used to derive an empirical distribution of the difference between F_exp_ and F_obs_.

### Selection scans in ancient populations

Our^27,28^ and published data (AADR) were used for imputation of the *IL23R* region. Only samples with genome-wide mean depth above 0.5X and breadth of coverage above 50%^29^ were considered for this part of the analysis, resulting in a total number of 151 individuals. The imputation was performed with GLIMPSE2^30^ using the 1000 Genomes Project reference panel^51^. To assess the accuracy of the imputation, we selected 24 high-coverage ancient genomes (>20X) for validation. Genotype calls for these genomes were generated using BCFtools v1.12^69^ with the same parameters described in the section above. BAM files for the high-coverage genomes were downsampled to 0.5X coverage with samtools and imputed with GLIMPSE2^30^ as previously described. We included positions that are in the 1000 Genomes Project strict accessibility mask, while filtering out high-repeat regions in RepeatMasker. The concordance of genotypes between validation and downsampled imputed datasets was done with GLIMPSE2_concordance with the following parameters: “--min-tar-gp --gt-val”. The bcftools command *consensus* was used to generate fasta files for the *IL23R*-phased haplotypes of each individual.

### Selection tests in ancient farmers and CEU

To test for significant selection signals around rs11209026, we applied various LD and site-frequency based methods. The LD based extended haplotype homozygosity (EHH) analysis can be used to detect recent positive selection^70^, even in ancient DNA datasets^71^. Haplotypes carrying positively selected variants are more extended compared to the ones under neutral conditions. EHH tests were conducted using the R package “rehh” version 3.2.2 with polarized VCFs that have diploid allele information^58^. By applying HWE, we tested whether the observed frequencies of heterozygotes and homozygotes for the derived allele aligned with the expectations under neutral (no selection) expectations. HWE and population differentiation (F_ST_) were calculated using PLINK version 2.0^67,68^. We further performed classic and haplotype-specific Tajima’s D, Fu and Li’s F, Fu and Li’s D tests that are site-frequency based methods^31,32^. Significant negative scores indicate positive selection or population expansion. The statistics were estimated by DnaSP 6^72^. The selection test could theoretically be affected by a bias in ancient DNA data. Both imputation and haplotype-specific analyses might reduce the number of singletons and rare variants, leading to more positive Tajima’s D etc. test results. This means that values calculated using biased ancient DNA data will be less statistically significant than those without any bias. In addition, the bias will affect both the selected and not selected haplotypes.

## Declarations

### Availability of data and materials

BAM files of aligned reads can be found at the European Nucleotide Archive (project accession no. XXXX). All other data needed to evaluate the findings in the paper are present in the main text or the supplementary materials.

